# Optogenetically Induced Microtubule Acetylation Unveils the Molecular Dynamics of Actin-Microtubule Crosstalk in Directed Cell Migration

**DOI:** 10.1101/2024.12.01.626286

**Authors:** Abhijit Deb Roy, Cristian Saez Gonzalez, Farid Shahid, Eesha Yadav, Takanari Inoue

## Abstract

Microtubule acetylation is implicated in regulating cell motility, yet its physiological role in directional migration and the underlying molecular mechanisms have remained unclear. This knowledge gap has persisted primarily due to a lack of tools capable of rapidly manipulating microtubule acetylation in actively migrating cells. To overcome this limitation and elucidate the causal relationship between microtubule acetylation and cell migration, we developed a novel optogenetic actuator, optoTAT, which enables precise and rapid induction of microtubule acetylation within minutes in live cells. Using optoTAT, we observed striking and rapid responses at both molecular and cellular level. First, microtubule acetylation triggers release of the RhoA activator GEF-H1 from sequestration on microtubules. This release subsequently enhances actomyosin contractility and drives focal adhesion maturation. These subcellular processes collectively promote sustained directional cell migration. Our findings position GEF-H1 as a critical molecular responder to microtubule acetylation in the regulation of directed cell migration, revealing a dynamic crosstalk between the actin and microtubule cytoskeletal networks.

## Introduction

Microtubules undergo at least nine different types of post-translational modifications, which independently, or in concert, modulate microtubule properties including its dynamics as well as their interaction with microtubule-associated proteins^1^. Acetylation of the lysine-40 residue of α-tubulin^2–6^, hereafter called microtubule acetylation for simplicity, is conserved throughout eukaryotes^7–9^, and is one of the only few modifications known to take place inside microtubule lumen^9^. Despite little significant structural changes^10^, microtubule acetylation provides structural stability against bending forces^11–16^. Microtubule acetylation has been implicated in cellular processes, including mechanosensing^12,17–21^, adaptation to extracellular environment^22–25^, intracellular transport via motor proteins^26–31^, DNA damage response^32^, autophagy^33–35^, and regulation of cell motility^22,23,36–39^. Directionally persistent cell migration, a process critical for physiological functions as well as pathological events, heavily relies on microtubule dynamics. While the involvement of microtubules in migration is well-established, the specific contributions of microtubule post-translational modifications remain poorly understood^40,41^. Microtubule acetylation, in particular, appears to exhibit differential roles in cell migration depending on cell type and environmental context. For example, it inhibits three-dimensional migration in human foreskin fibroblasts^23^ and transwell migration in NIH3T3 fibroblasts^42^ while promoting motility in astrocytes^22,24^ and breast cancer cells^36–38^. In contrast, acetylation is dispensable for the motility of RPE1 epithelial cells^43^. These conclusions have largely been drawn from studies employing genetic engineering or pharmacological interventions to alter microtubule acetylation.

Genetic approaches are invaluable for identifying genes responsible for these effects; however, they often lack the temporal precision required to investigate rapid cytoskeletal dynamics during cell migration, which can occur within minutes. Similarly, pharmacological interventions to modulate microtubule acetylation may have unintended non-specific effects^42,44–47^, complicating the interpretation of results. The paucity of molecular tools capable of controlling microtubule acetylation with rapid temporal resolution and high molecular specificity has presented a significant challenge in elucidating its real-time roles in dynamic cell behavior such as directional cell migration. To address this, we developed a genetically encoded actuator, termed optoTAT, that is designed to induce microtubule acetylation within minutes upon light illumination. By leveraging this optogenetic actuator in combination with genetic knock-out models, migration assays, and live cell fluorescence imaging, we aimed to uncover the molecular interplay between microtubule acetylation, actin cytoskeleton remodeling, and directional cell migration in real time.

## Results

### Microtubule acetylation mediates directional migration

α-TAT1 is the only enzyme known to acetylate microtubules in mammals^12,48,49^, whereas the deacetylation is catalyzed by HDAC6 and Sirt2^42,50^. Mouse Embryonic Fibroblasts (MEFs) obtained from α-TAT1 knockout (KO) mice do not have detectable microtubule acetylation^12,17,51^. In a random migration assay, α-TAT1 KO MEFs showed significantly greater motility but reduced directional persistence compared to wild-type (WT) MEFs (Fig. 1a, b, c). In wound healing assays, α-TAT1 KO MEFs closed the wound more rapidly than WT MEFs (Fig.1d, e). To examine the effects of microtubule acetylation on chemotaxis, we utilized an Ibidi chemotaxis chamber with 0-20% FBS gradient (Fig. 1f). α-TAT1 KO MEFs failed to efficiently migrate towards the chemoattractant compared to WT MEFs (Fig. 1g, Supplementary Fig. S1a). Unlike the WT MEFs, the α-TAT1 KO MEFs exhibited reduced directional bias towards the chemoattractant gradient, as indicated by the reduced shift in the center of mass from the origin (Supplementary Fig. S1b). The α-TAT1 KO MEFs showed significantly reduced directional persistence compared to WT MEFs, as shown by decreased forward migration index (FMI) for the KO MEFs along the chemoattractant gradient (FMI^II^) (Fig. 1h), but not perpendicular to the gradient (FMI^⊥^) (Fig. 1i). These chemotaxis defects were rescued by exogenous expression of mVenus-α-TAT1 in KO MEFs (Fig. 1g, h, Supplementary Fig. S1a, b).

**Figure 1.**
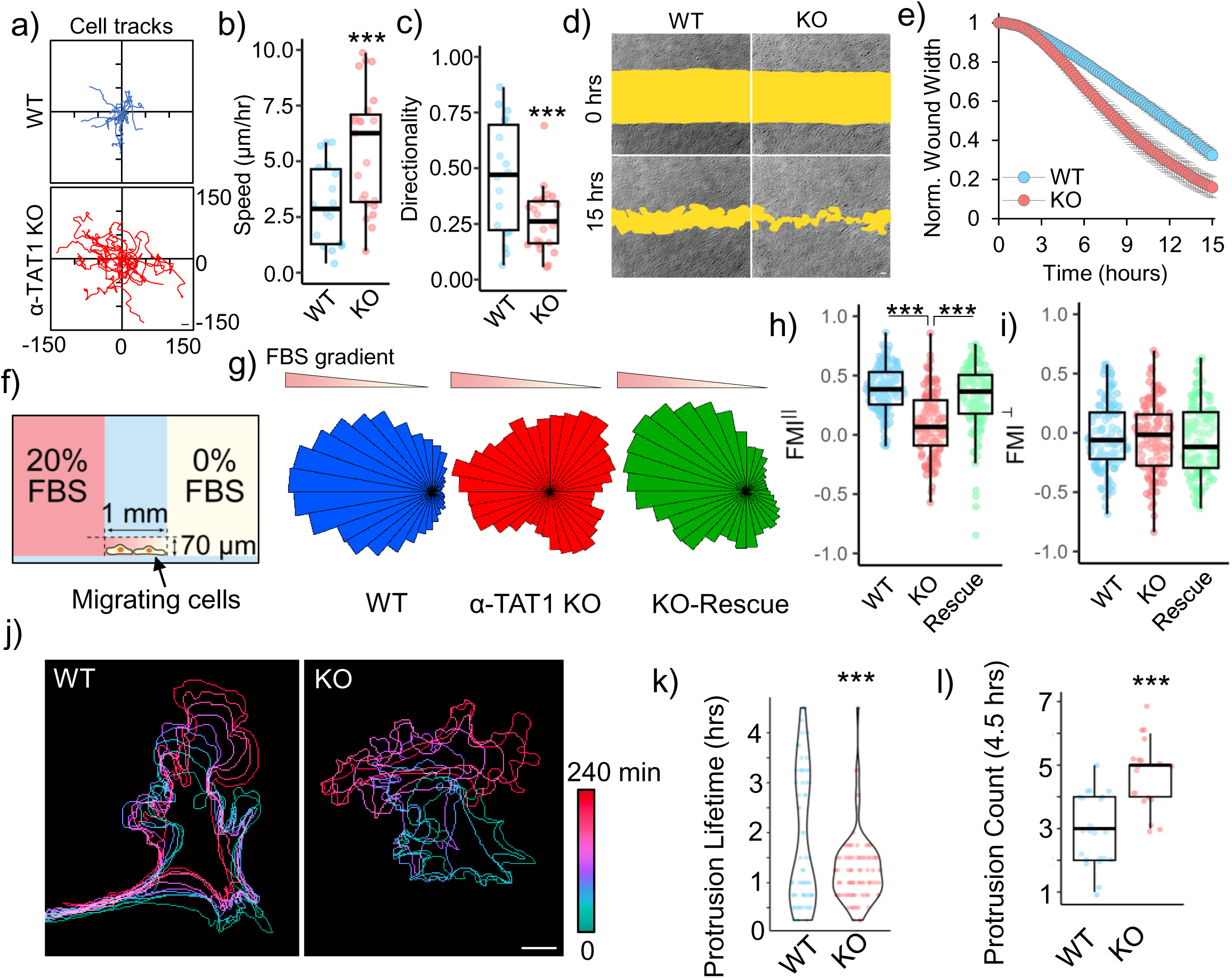
α-TAT1 modulates directional cell migration. a) Tracks, b) Speed (µm/hr) and c) Directionality of WT and α-TAT1 KO MEFs in a random migration assay, WT: 18, KO: 23 cells, scale bar: 10 μm; d), e) Temporal changes in wound width in a wound healing assay with WT and α-TAT1 KO MEFs, n = 12 wound regions from 3 independent experiments, mean ± 95% C.I.; f) Schematic for chemotaxis assay *(adapted from Ibidi);* g) Rose plots of WT, α-TAT1 KO or KO-rescue MEFs migrating in a chemotactic gradient, h) Forward migration indices along the chemotactic gradient and i) Forward migration indices perpendicular to the chemotactic gradient for WT, α-TAT1 KO or KO-rescue MEFs, n = 120 cells (40 each from three independent experiments); j) Temporal changes in morphology of WT or α-TAT1 KO MEFs undergoing random migration, scale bar: 10 μm; k) Persistence of protrusions, l) Frequency of new protrusion formation in randomly migrating WT or α-TAT1 KO MEFs, WT: 23 and KO: 19 cells. ***: p<0.001

On examining the motility of WT and α-TAT1 KO MEFs, we observed that in contrast to WT MEFs, the α-TAT1 KO MEFs change their direction of motion repeatedly (Fig. 1j). The WT MEFs had two groups of protrusions, one short-lived and another long-lived, as indicated by the bimodal distribution of protrusion lifetimes. α-TAT1 KO MEFs had very few such long-lived protrusions (Fig. 1k). The α-TAT1 KO MEFs also produced new protrusions more frequently than WT MEFs, leading to changes in direction of movement (Fig. 1l). Directional persistence requires a long-lasting front-back polarity^52,53^, and the frequent protrusion formation in α-TAT1 KO MEFs suggest defects in maintenance of such front-back polarity. Consistent with this, morphological analyses showed that α-TAT1 KO MEFs have higher circularity and higher convexity (Supplementary Fig. S1c, d), consistent with more protrusive phenotypes. The chemotaxis defects in α-TAT1 KO MEFs were not due to defects in sensing the chemoattractant since serum-starved WT and α-TAT1 KO MEFs showed comparable morphological changes: increased protrusions, on treatment with 10% FBS (Supplementary Fig. S1e).

### Microtubule acetylation mediates focal adhesion maturation

In migrating cells, nascent protrusions are stabilized by integrin mediated adhesion complexes which undergo maturation in response to actomyosin contractility^54,55^. On immunostaining for Vinculin, we observed fewer adhesions in α-TAT1 KO MEFs compared to WT MEFs (Fig. 2a, b). Consistent with decreased adhesion, α-TAT1 KO MEFs also had smaller cell spread area (Supplementary Fig. S2a). α-TAT1 KO MEFs also had fewer larger mature adhesions (Fig. 2a, Supplementary Fig. S2b), suggesting a defect in the adhesion maturation pathways. In WT MEFs, we observed a polarized distribution of nascent and maturing adhesions in the front, and large mature adhesions at retractions, whereas the α-TAT1 KO MEFs lacked any such spatial polarization of adhesions (Fig. 2a). We also observed lower levels of vinculin localization in the adhesions in the α-TAT1 KO MEFs (Fig. 2a, Supplementary Fig. S2c, d), which is consistent with lower tensile forces actin on these adhesions. Exogenous expression of mVenus-α-TAT1, but not a catalytically dead mVenus-α-TAT1(D157N)^12^, in α-TAT1 KO MEFs could rescue these adhesion defects (Fig.2a, b, Supplementary Fig. S2b, c, d). These adhesion defects were not due to decreased Vinculin expression since both WT and α-TAT1 KO MEFs showed comparable levels of Vinculin expression (Fig. 2c, d). Adhesion maturation is mediated by tensile forces experienced by focal adhesion components through the actin cytoskeleton. Using a Vin-TS FRET-based tension sensor^56^ we observed increased FRET in the α-TAT1 KO MEFs, indicating that focal adhesions experienced significantly reduced forces compared to WT MEFs (Fig. 2e, f, Supplementary Fig. S2e). Furthermore, the Vin-TS FRET signal also showed a polarized distribution in the WT MEFs, indicating a polarized distribution of tensile forces, but not in the KO cells (Fig. 2e).

**Figure 2.**
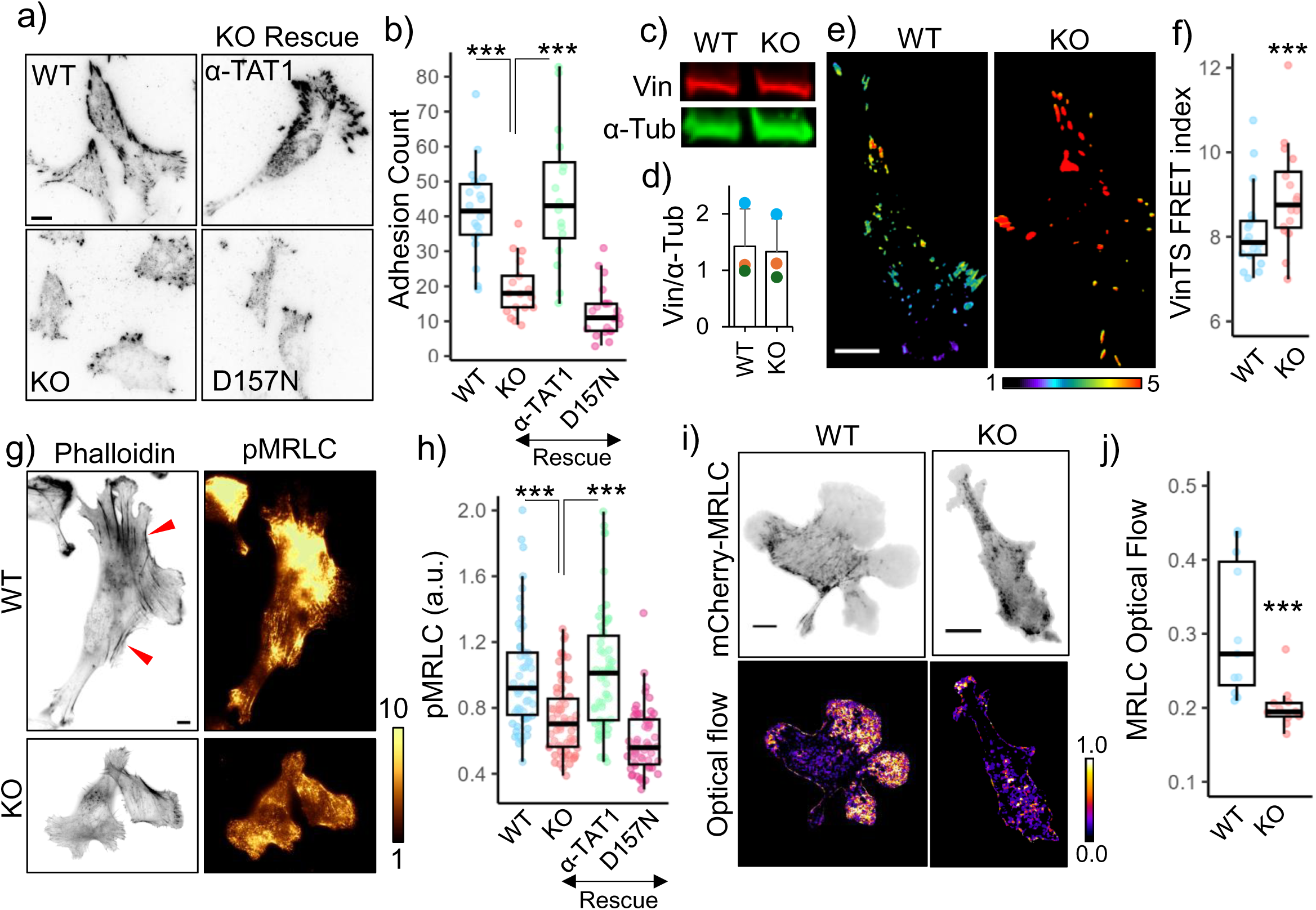
Microtubule acetylation promotes focal adhesion maturation and actomyosin contractility. a) Vinculin distribution in WT, α-TAT1 KO MEFs, and KO-rescue with mVenus-α-TAT1 or catalytic dead mVenus-α-TAT1(D157N) as indicated; b) Number of adhesions per cell (WT:20, KO: 17, rescue-WT: 16, rescue-D157N: 22 cells); c) Western blot showing Vinculin and α-Tubulin expression in WT and α-TAT1 KO MEFs; d) Normalized Vinculin expression levels in WT and α-TAT1 KO MEFs by Western blots (3 independent experiments, error bar: standard deviation); e) VinTS FRET index in WT and α-TAT1 KO MEFs, f) Average VinTS FRET index in WT and α-TAT1 KO MEFs (WT:: 18, KO: 16 cells); g) Phalloidin and phospho-MRLC distribution in WT and α-TAT1 KO MEFs, red arrowheads indicate bundled actin; h) Phospho-MRLC levels in WT, α-TAT1 KO, rescue-WT and rescue-D157N MEFs (WT: 54, KO: 64, rescue-WT: 53 and rescue-D157N: 55 cells); i) mCherry-MRLC distribution and optical flow levels of mCherry-MRLC in WT and α-TAT1 KO MEFs; j) Mean mCherry-MRLC optical flow levels in WT and α-TAT1 KO MEFs (WT: 11, KO: 12 cells). Scale bar: 10 μm. ***: p<0.001

### Microtubule acetylation mediates actomyosin contractility

Focal adhesions experience tensile forces through the actin cytoskeleton^57^. Phalloidin staining showed a significant reduction in bundled actin in α-TAT1 KO cells, suggesting that these cells have defects in actin contractility (Fig. 2g, red arrowheads). Contractility in the actin cytoskeleton is generated through Myosin motor proteins, which are activated through phosphorylation of the Myosin Regulatory Light Chain (MRLC) at Serine19 by Myosin Light Chain Kinase (MLCK)^58^. Myosin activation is also involved in directional persistence of migrating cells^59^. Immunostaining of WT or α-TAT1 KO MEFs with an antibody against phospho-MRLC Serine19 showed a significantly lower levels of phospho-MRLC (Fig. 2g, h), indicating decreased activation levels of Myosin. The decrease in phospho-MRLC levels was not due to a decrease in expression levels since WT and KO cells showed comparable Myosin expression levels (Supplementary Fig. S2f, g). These defects in MRLC phosphorylation could be rescued with exogenous expression of mVenus-α-TAT1 but not mVenus-α-TAT1(D157N) mutant (Fig. 2h). Since MRLC phosphorylation leads to association with the actin cytoskeleton, we measured optical flow of mCherry-MRLC to characterize myosin activation dynamics. mCherry-MRLC flow was considerably lower in α-TAT1 KO MEFs compared to WT cells (Fig. 2i, j), indicating decreased association with the actin cytoskeleton. Treatment of KO cells with Y-27632 led to a decrease in phospho-Myosin levels, suggesting a residual amount of myosin activity, however diminutive (Supplementary Fig. S2h, i).

### Inhibiting HDAC6 weakly promotes myosin activation

Our observations suggest that microtubule acetylation promotes myosin activation. Tubacin is a widely used pharmacological inhibitor of HDAC6^60^. To test whether increase in microtubule acetylation levels could increase myosin activation, we used TIRF microscopy to characterize changes in mCherry-MRLC association with actin cytoskeleton in WT MEFs treated with 2 μM Tubacin. Over 5 hours post Tubacin treatment, we observed a minor increase in mCherry-MRLC signal on the TIRF plane (Supplementary Fig. S2j, k), indicating increased activation and association with the actin cytoskeleton. However, this increase over the course of hours does not eliminate the possibility of cell adaptation through transcriptional regulation or non-specific effects. HDAC6 has many other substrates other than α-Tubulin^44,61,62^, and it can also deacetylates additional acetylated lysine residues on α-Tubulin^63^. Thus, HDAC6 inhibition is not sufficiently specific to determine causal relationships between acetylation of Lysine-40 in α-Tubulin and cellular or molecular responses.

### Developing an optogenetic actuator to rapidly induce microtubule acetylation

To examine a specific and causal relationship between microtubule acetylation and myosin activation, we sought to develop an inducible molecular actuator to control microtubule acetylation. Initially we tested Z-lock-α-TAT1 in HeLa cells^64^. However, we observed a significant increase in microtubule acetylation in cells expressing mCherry-Z-lock-α-TAT1 even in dark (Supplementary Fig. 3a). We have previously shown that cytoplasmic localization of α-TAT1 through its C-terminal spatial regulatory domain is critical for microtubule acetylation. Nuclear localization of α-TAT1 is sufficient to sequester it from catalyzing microtubule acetylation^51^. Based on this, we reasoned that inducing export of a nuclear-localized α-TAT1 may induce acetylation of microtubules (Fig. 3a). We initially implemented the light-inducible nuclear export system (LEXY)^65^ to sequester full-length α-TAT1(M1-R323) in the nucleus in dark. We named this construct <u>Opto</u>genetic <u>T</u>ubulin <u>A</u>cetyl-<u>T</u>ransferase <u>v</u>ersion <u>0</u> (optoTATv0) (Fig. 3b). On blue-light stimulation, we observed a rapid nucleus-to-cytoplasm translocation of mCherry-optoTATv0 (Fig. 3c, d). However, we also observed significant levels of cytoplasmic presence even in the absence of blue-light stimulation (Fig. 3c, d), presumably due to the presence of nuclear export and cytoplasmic retention machinery in α-TAT1 C-terminus^51^. To improve upon this design, we tethered only the catalytic domain of α-TAT1(M1-S236)^12^ to LEXY (optoTATv1) or to LEXY with two NLS (optoTATv2) (Fig. 3b). These versions showed increased nuclear sequestration (Fig. 3c, d), with rapid, robust and reversible cytoplasmic translocation on blue light stimulation (Fig. 3c, d, e, f,). To examine whether blue-light stimulation of these tools could acetylate microtubules, we exposed HeLa cells expressing mCherry-optoTATv1 or mCherry-optoTATv2 to blue light for 2 hours and performed immunostaining for acetylated microtubules. We observed that blue light stimulation of optoTATv1 or optoTATv2 significantly induced microtubule acetylation in HeLa cells (Fig. S3g, h). However, we observed that the cells expressing optoTATv1, but not optoTATv2, showed increased levels of microtubule acetylation in dark when compared to non-transfected cells (Fig. 3g, h). We used optoTATv2 for all further experiments, and for simplicity, we will refer to it as optoTAT here onwards.

**Figure 3.**
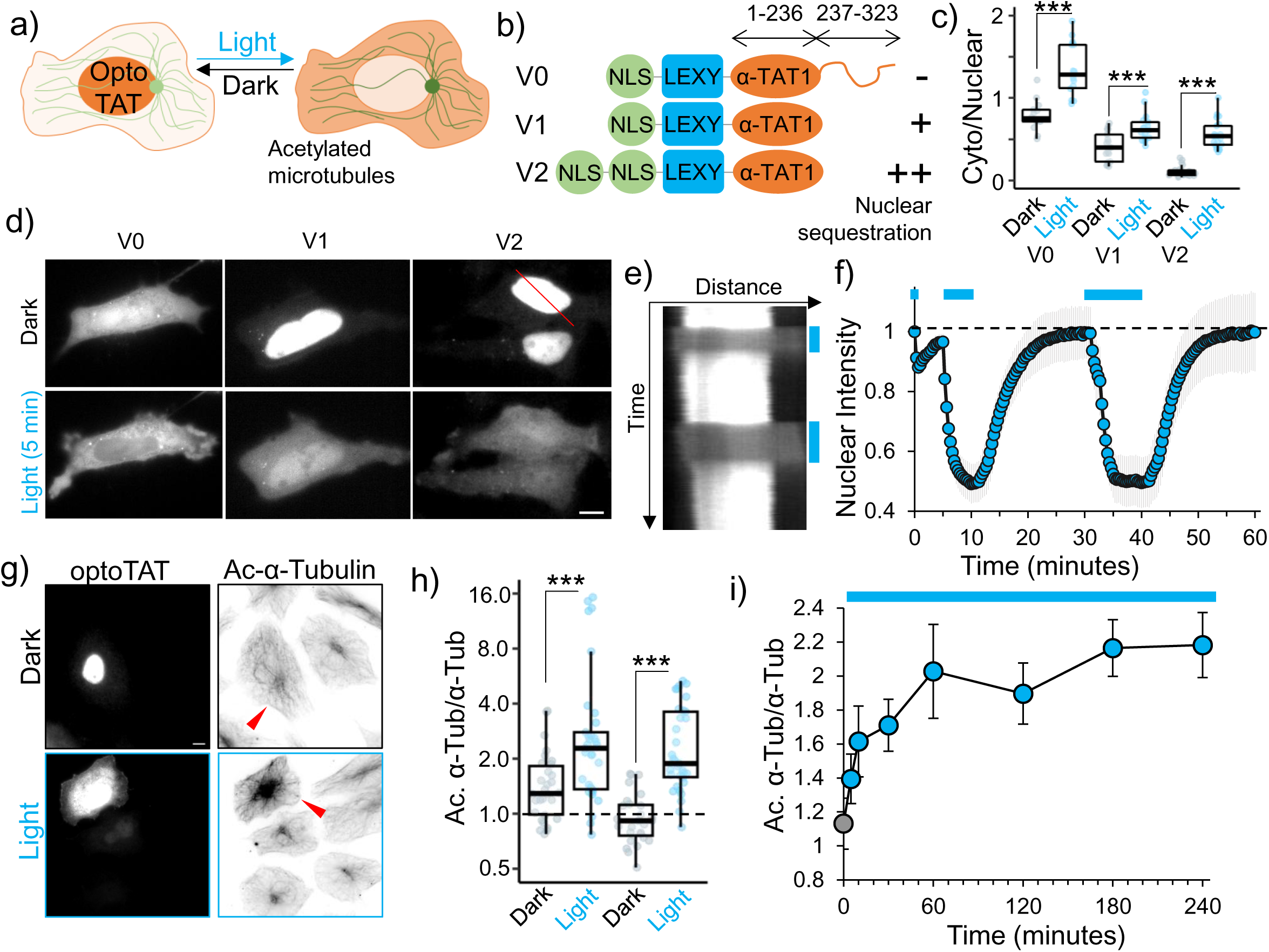
Developing an optogenetic actuator to induce microtubule acetylation. a) OptoTAT design principle; b) OptoTAT versions; c) Ratio of cytoplasmic over nuclear signal for different versions of optoTAT in dark and on 10 min blue light stimulation (V0: 14, V1: 17 and V2: 21 cells); d) Changes in intracellular distribution of mCherry-optoTAT V0, V1 and V2 in dark and on blue light stimulation; e) Kymograph showing mCherry-optoTAT V2 response to blue light, reference for kymograph is the red line in top panel of (d); f) Temporal changes in average nuclear intensity of mCherry-optoTAT V2 on blue light stimulation indicated by blue lines, means ± 95% C.I. are shown, n= 21 cells; g) Microtubule acetylation levels in HeLa cells exogenously expressing mCherry-optoTAT V2, kept in dark or exposed to blue light for 2 hours, red arrowheads indicate transfected cells; h) Acetylated microtubule levels (normalized against total α-Tubulin) in HeLa cells expressing mCherry-optoTAT V1 or V2 in dark or exposed to 2 hours blue light, values were normalized against non-transfected cells in the same dish (V1 dark: 30, V1 light: 34, V2 dark: 27, V2 light: 33 cells); i) Temporal changes in levels of acetylated microtubules (normalized against total α-Tubulin) in HeLa cells stably expressing mVenus-optoTAT and continuously exposed to blue light stimulation for the duration indicated (0 min: 54, 5 min: 50, 10 min: 61, 30 min: 66, 60 min: 61, 120 min: 62, 180 min: 60 and 240 min: 61 cells), means ± 95% C.I. are shown. Scale bar: 10 μm.

To assess the kinetics of microtubule acetylation by optoTAT, we used lentiviral transduction to generate a cell line of HeLa cells stably expressing mVenus-optoTAT. We used flow cytometric cell sorting to select cells with comparable levels of mVenus expression. These cells were incubated in dark for 24 hours and then exposed to blue-light for 0 min, 5 min, 10 min, 30 min, 60 min, 120 min and 240 min, followed by immunostaining for acetylated α-Tubulin and total α-Tubulin. We observed a rapid and significant increase in microtubule acetylation within 10 minutes of blue-light stimulation, which continued to increase and stabilize after an hour of stimulation (Fig. 3i, Supplementary Fig. S3b,c,d). Our data demonstrate that we have developed an optogenetic molecular actuator to rapidly induce microtubule acetylation in living cells, with a tunable dynamic range.

### OptoTAT stimulation rapidly induces myosin activation

Phosphorylation of MRLC at Serine-19, leads to Myosin (and MRLC) activation, resulting in association with F-actin^58^. We stimulated miRFP703-optoTAT in HeLa cells co-expressing mCherry-MRLC and visualized changes in MRLC distribution using TIRF microscopy. We reasoned that Myosin activation will coincide with an increased MRLC association with the actin cytoskeleton, leading to an increase in mCherry-MRLC fluorescence in the TIRF plane^66^. On blue light stimulation, we observed a rapid and persistent increase in mCherry intensity in the TIRF plane, indicating increased association of mCherry-MRLC with actin cytoskeleton (Fig. 4a, b,). This increase was concurrent with increased coherence in mCherry-mRLC distribution (Fig. 4c, d), suggesting increased isotropy in myosin distribution, consistent with higher levels of bundled actin and increased actomyosin contractility. Catalytically dead miRFP-optoTAT(D157N) failed to elicit any increase in mCherry-MRLC signal. Additionally, any increase in mCherry-MRLC intensity was abrogated on treating the cells with ROCK inhibitor Y27632 for 10 minutes before optoTAT stimulation, or by pre-saturating microtubule acetylation by treating the cells with HDAC6 inhibitor tubacin for 4 hours before optoTAT stimulation (Fig. 4e).

**Figure 4.**
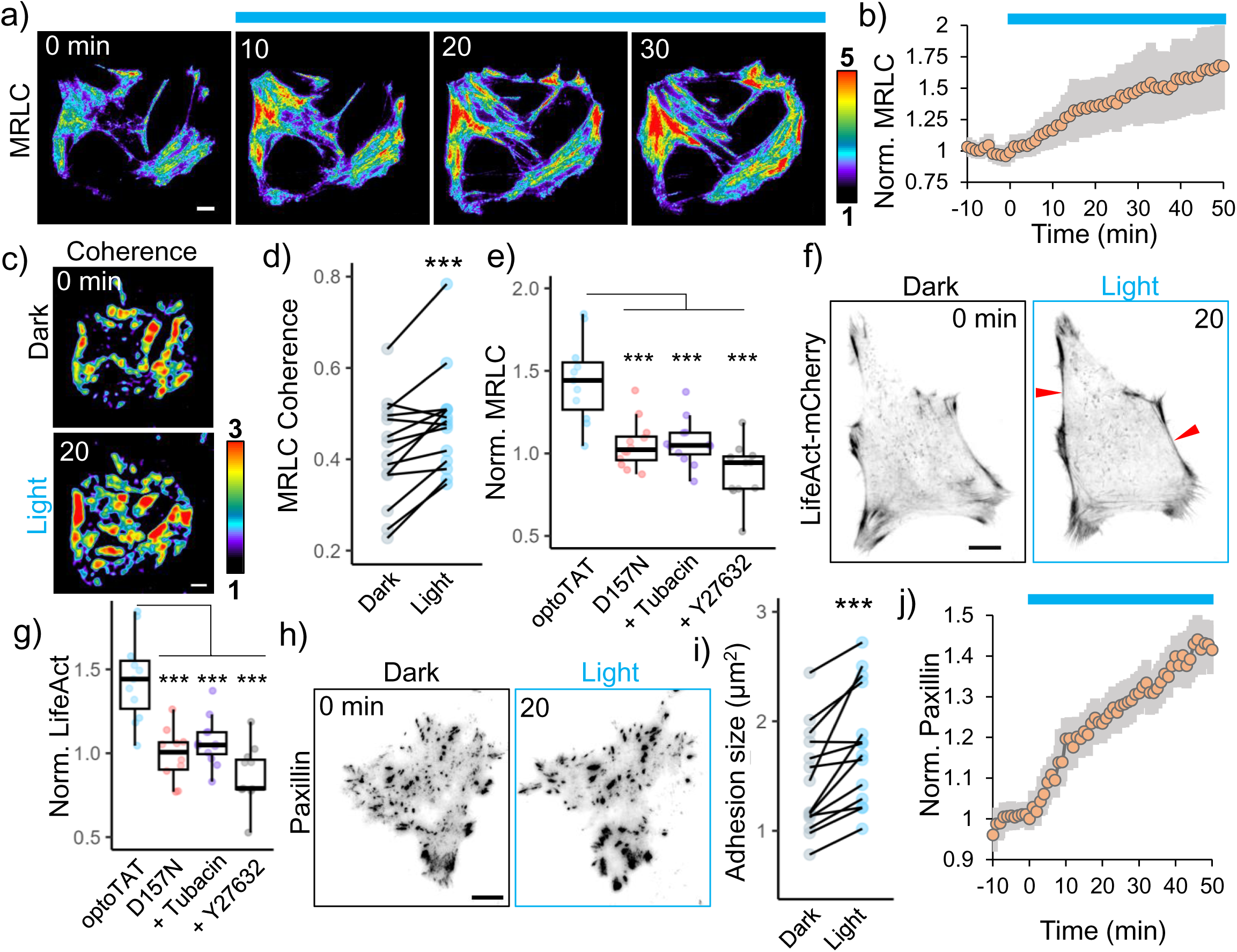
OptoTAT stimulation rapidly induces actomyosin contractility. a) TIRF images showing temporal changes in mCherry-MRLC distribution on miRFP703-optoTAT stimulation in HeLa cells; b) Changes in mCherry-MRLC intensity on miRFP703-optoTAT stimulation, mean ± 95% C.I. are shown, n = 14 cells, c), d) Changes in MRLC distribution isotropy on miRFP703-optoTAT stimulation, n = 14 cells; e) Changes in mCherry-MRLC intensity in TIRF plane on 30 min blue light stimulation of miRFP703-optoTAT (14 cells), catalytically dead miRFP703-optoTAT(D157N) (12 cells), miRFP703-optoTAT and pre-treatment with 2 µM Tubacin (12 cells) or 10 µM Y27632 (12 cells); f) Changes in LifeAct-mCherry on miRFP703-optoTAT stimulation, red arrowheads indicate bundled actin; g) Changes in LifeAct-mCherry intensity in TIRF plane on 30 min blue light stimulation of miRFP703-optoTAT (12 cells), catalytically dead miRFP703-optoTAT(D157N) (12 cells), miRFP703-optoTAT and pre-treatment with 2 µM Tubacin (12 cells) or 10 µM Y27632 (10 cells), h) Changes in mCherry-Paxillin on miRFP703-optoTAT stimulation, i) Changes in average focal adhesion sizes and j) changes in average mCherry-Paxillin intensity on 30 min miRFP703-optoTAT stimulation, mean ± 95% C.I. are shown, n = 14 cells. Scale bar: 10 μm. ***: p<0.001. Blue line: Blue light stimulation.

Consistent with an increase in myosin activity, miRFP703-optoTAT stimulation also led to increased levels of bundled actin (Fig. 4f, g), and maturation of focal adhesions as indicated by increased adhesion sizes and mCherry-Paxillin accumulation (Fig. 4h, i, j,). Taken together, these data suggest that optoTAT stimulation rapidly induced Myosin activation through increased microtubule acetylation and ROCK kinase activation.

### Microtubule acetylation releases GEF-H1 from sequestration

Myosin activation on MRLC phosphorylation through MLCK is often downstream of RhoA-ROCK signaling. Our observation that inhibiting ROCK abrogated optoTAT mediated Myosin activation suggested that optoTAT stimulation leads to RhoA activation. GEF-H1 is an activator for RhoA, which is sequestered on microtubules and is activated on disrupting microtubules using nocodazole^67,68^. GEF-H1 was reported to mediate α-TAT1 mediated cellular mechano-sensing^22,69^. Immunostaining revealed significantly increased GEF-H1 sequestration on the microtubules in α-TAT1 KO cells compared to WT cells (Fig. 5a, b). This was not due to increased expression levels since both WT and α-TAT1 KO MEFs had comparable GEF-H1 expression levels (Fig. 5c, d). Since acetylation has been reported to stabilize microtubules, we speculated whether release of GEF-H1 was due to increased stability of microtubules in WT MEFs. However, we did not detect any significant loss of GEF-H1 sequestration in α-TAT1 KO cells on treatment with Paclitaxel (Supplementary Fig. S4a), suggesting that an increase in microtubule stability, or protection from disassembly, did not significantly affect GEF-H1 sequestration. To examine if the acetyl moiety specifically was responsible for GEF-H1 release, we co-immunostained for GEF-H1, acetylated microtubules and α-Tubulin in WT MEFs that were treated with Paclitaxel to eliminate any potential effects of microtubule stability. We observed a negative correlation between the spatial distribution of microtubule-bound GEF-H1 and acetylated microtubules (Fig. 5e, f, Supplementary Fig.S4b). To test whether GEF-H1 specifically binds to non-acetylated microtubules, we exogenously expressed mCherry-α-Tubulin or acetylation deficient mCherry-α-Tubulin(K40A) mutant in HeLa cells and immunostained for GEF-H1. We observed increased microtubule-bound GEF-H1 in the cells expressing mCherry-α-Tubulin(K40A), compared to non-transfected cells, or those expressing WT mCherry-α-Tubulin (Supplementary Fig. S4c).

**Figure 5.**
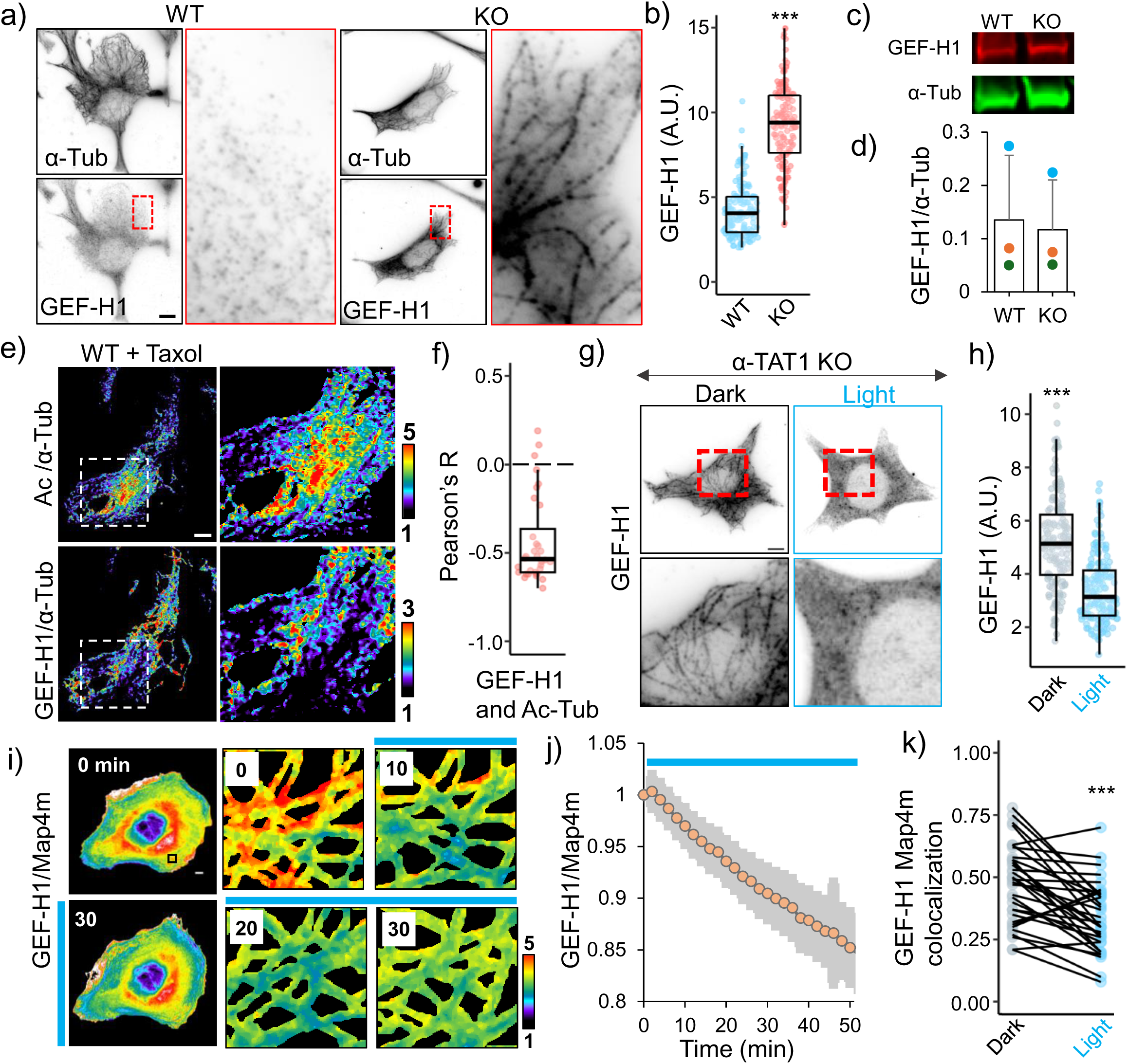
Microtubule acetylation releases GEF-H1 sequestration. a) α-Tubulin and GEF-H1 localization in WT and α-TAT1 KO MEFs, inset for GEF-H1 is magnified on right; b) linear density of GEF-H1 along microtubules in WT and α-TAT1 KO MEFs (5 microtubules from 30 cells each, total 150); c), d) GEF-H1 expression levels in WT and α-TAT1 KO MEFs measured using Western blots (3 independent experiments, error bar: standard deviation); e) Relative distributions of acetylated microtubules (top panel) and microtubule-bound GEF-H1 (bottom panel) in overnight 100 nM Taxol treated WT MEFs, inset magnified on the right panels; f) Pearson’s R value for spatial colocalization of acetylated microtubules and microtubule-bound GEF-H1 in Taxol treated WT MEFs, n = 33 cells; g) GEF-H1 localization in α-TAT1 KO MEFs stably expressing mVenus-optoTAT kept in dark or with 30 min blue light stimulation; h) linear density of GEF-H1 along microtubules in α-TAT1 KO MEFs stably expressing mVenus-optoTAT kept in dark or exposed to 30 min blue light stimulation (5 microtubules from 30 cells each, total 150); i) Changes in mCherry-GEF-H1/mVenus-MAP4m signal in HeLa cells expressing miRFP703-optoTAT on blue light stimulation, inset is magnified in the right panels; j) Temporal changes in mCherry-GEF-H1/mVenus-MAP4m on miRFP703-optoTAT stimulation, mean ± 95% C.I. are shown, n = 33 cells; k) Changes in colocalization of mCherry-GEF-H1 and mVenus-Map4m on miRFP703-optoTAT stimulation for 30 min, n = 33 cells. Scale bar: 10 μm. ***: p<0.001

On stimulating α-TAT1 KO MEFs expressing mVenus-optoTAT with blue light and immunostaining for GEF-H1, we observed reduced localization of endogenous GEF-H1 on microtubules compared to those the cells kept in dark. (Fig. 5g, h). To examine the release kinetics of GEF-H1 on microtubule acetylation, we exogenously expressed mCherry-GEF-H1, EYFP-Map4m (Map4 microtubule binding domain) and miRFP703-optoTAT in HeLa cells and characterized changes in mCherry-GEF-H1 distribution on blue light stimulation. Since YFP excitation was sufficient to activate optoTAT, we could not obtain any ‘before’ images except time zero. Nevertheless, we observed a persistent decrease in mCherry/EYFP signal, indicating a release of GEF-H1 from microtubules (Fig. 5i, j, Supplementary Fig. S4d,). Cross-correlation analysis of mCherry and EYFP signal in dark or after 20 min stimulation also showed a decrease, further confirming release of GEF-H1 (Fig. 5k). These data suggest that microtubule acetylation rapidly releases sequestered GEF-H1 to activate RhoA and actomyosin contractility.

### GEF-H1 release mediates microtubule acetylation dependent myosin activation

To test whether GEF-H1 mediated optoTAT induced myosin activation, we used RNAi to deplete GEF-H1 in HeLa cells (Fig. 6a, b) and examined changes in mCherry-MRLC signal on miRFP703-optoTAT stimulation. Cells treated with siRNA against GEF-H1, but not the control siRNA, did not show significant increase in mCherry-MRLC signal on optoTAT activation (Fig. 6c). To examine whether release of GEF-H1 from microtubules was critical for microtubule acetylation mediated myosin activation, we used lentiviral transduction to stably express mCherry-GEF-H1(C53R) in α-TAT1 KO MEFs. GEF-H1(C53R) is a mutant that does not bind to microtubules (Fig. 6d) but retains its capability to activate RhoA^70^. mCherry-GEF-H1(C53R) expression was sufficient to increase phospho-MRLC levels in α-TAT1 KO MEFs compared to non-transduced ones in a ROCK kinase dependent manner (Fig. 6e, f). These data suggest that microtubule acetylation dependent activation of Myosin was mediated by GEF-H1.

**Figure 6.**
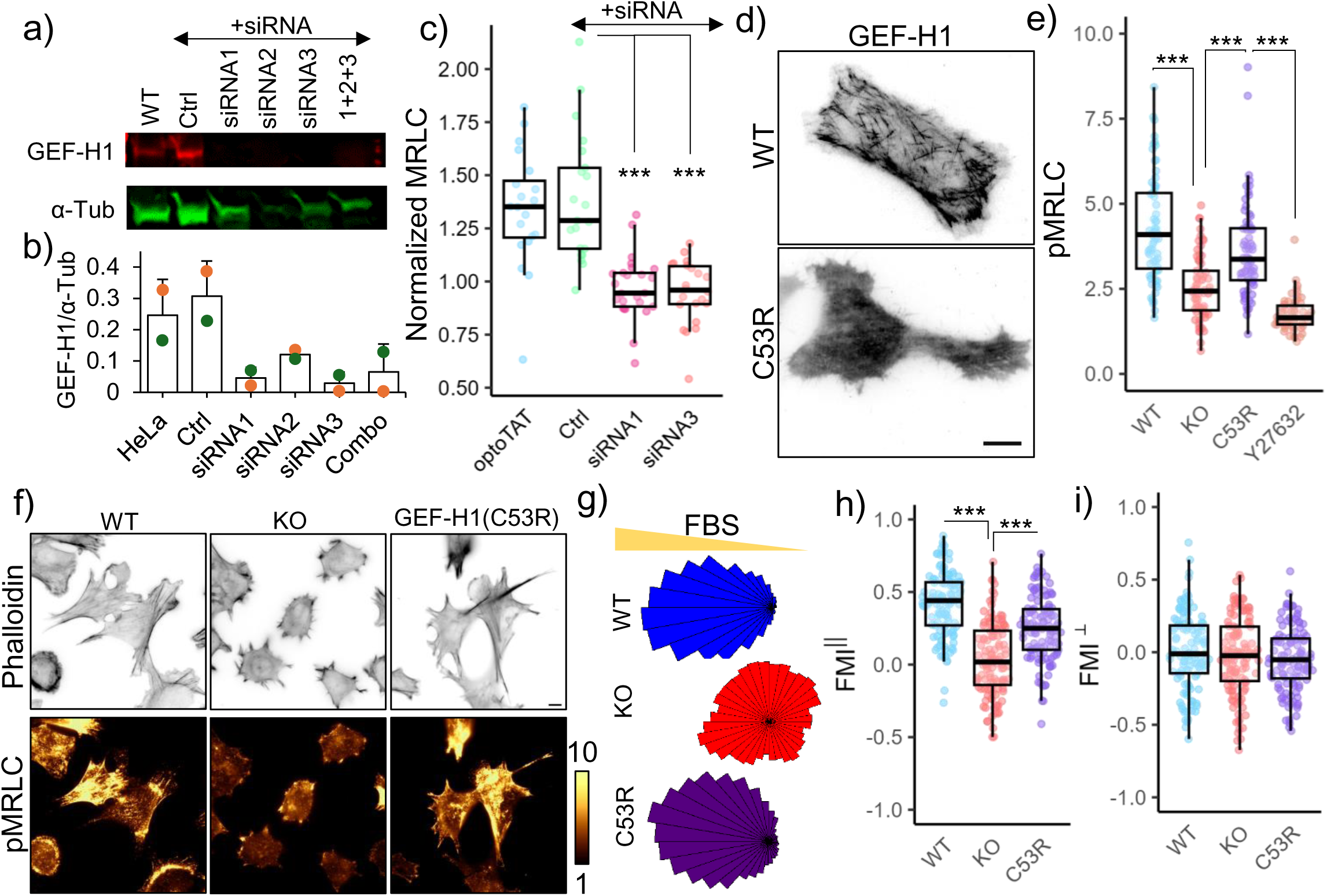
GEF-H1 mediates microtubule acetylation dependent actomyosin contractility. a), b) GEF-H1 knock-down in HeLa cells by siRNA; c) Changes in mCherry-MRLC intensity on miRFP703-optoTAT stimulation in HeLa cells with optoTAT (20 cells), scramble siRNA (21 cells), siRNA1 (25 cells) and siRNA3 (22 cells) against GEF-H1; d) TIRF images of HeLa cells expressing GFP-GEF-H1 (top panel) and mCherry-GEF-H1(C53R); e) Phospho-MRLC levels in WT (67 cells), α-TAT1 KO (69 cells), α-TAT1 KO MEFs expressing mCherry-GEF-H1(C53R) (67 cells) and same cells treated with 10 µM Y-27632 (60 cells); f) Phalloidin and phospho-MRLC distribution in WT, α-TAT1 KO MEFs and α-TAT1 KO MEFs expressing mCherry-GEF-H1(C53R); g) Rose plots of WT, α-TAT1 KO and KO-GEF-H1(C53R) MEFs migrating in a chemotactic gradient; g) Forward migration indices along the chemotactic gradient and h) Forward migration indices perpendicular to the chemotactic gradient for WT, α-TAT1 KO and KO-GEF-H1(C53R) MEFs, n = 120 cells (40 each from three independent experiments). Scale bar: 10 μm. ***: p<0.001

### Microtubule binding deficient GEF-H1 rescues chemotaxis defects of α-TAT1 KO MEFs

Based on our observations, we hypothesized that the defects in directional migration of α-TAT1 KO MEFs is due to defects in release of GEF-H1 from microtubules, resulting in lower actomyosin contractility. If this were true, expressing GEF-H1(C53R) in α-TAT1 KO MEFs should rescue their defects in chemotaxis. We performed chemotaxis assay with WT, α-TAT1 KO MEFs and α-TAT1 KO MEFs expressing mCherry-GEF-H1(C53R) in 0-20% FBS gradient. α-TAT1 KO-GEF-H1(C53R) MEFs showed significantly improved chemotactic capability compared to α-TAT1 KO MEFs, at levels comparable to WT MEFs (Fig. 6f, g, h, Supplementary Fig. S5a, b). Altogether, these data demonstrate that microtubule acetylation drives directional migration by modulating actomyosin contractility in migrating cells through dynamic release of sequestered GEF-H1.

## Discussion

Our data suggest that microtubule acetylation reduces overall motility in MEFs, but facilitates directional motility by promoting a dominant protrusion, whilst inhibiting nascent ones (Fig. 1). This coordination was achieved by modulation of actomyosin contractility and stabilizing adhesions in the dominant protrusion (Fig. 2). While microtubules have been implicated in focal adhesion turnover and actomyosin contractility through Rac1 and RhoA^71–77,68^, the specific effects of microtubule acetylation on myosin activation is unclear. In astrocytes and HUVEC cells, microtubule acetylation promotes myosin activation through GEF-H1, whereas in human foreskin fibroblasts microtubule acetylation inhibits myosin activity through MYPT1^22,23^. Our data demonstrate that migration defects in α-TAT1 KO MEFs arise from decreased actomyosin contractility through sequestration of GEF-H1 in non-acetylated microtubules (Fig. 6). Consistently, we observed rapid myosin activation on optoTAT stimulation through GEF-H1 release (Fig. 4, 5) in MEFs and HeLa cells. Intriguingly, microtubule acetylation mediated increased actomyosin contractility promoted astrocyte migration^22^, whereas decreased myosin activation in human foreskin fibroblasts due to microtubule acetylation inhibited migration^23^. Our data suggest an overall decrease in MEF motility due to increased actomyosin contractility (Fig. 1, 2). Although these variations in migratory phenotypes suggest a context dependent role of actomyosin contractility in migrating cells, our findings using optoTAT provide evidence for a specific molecular coupling between microtubule acetylation in GEF-H1 release and myosin activation. It should be noted that optoTAT lacks the C-terminus of α-TAT1, which may prevent it from fully recapitulating all signaling pathways involving α-TAT1. Additionally, the nuclear sequestration of optoTAT may influence cell behavior. Nevertheless, the absence of myosin activation upon stimulation with catalytically inactive optoTAT(D157N) (Fig. 4) and the increased microtubule sequestration of GEF-H1 in cells expressing α-Tubulin(K40A) (Supplementary Fig. S4) strongly support a specific role for microtubule acetylation in facilitating GEF-H1 release and myosin activation. We want to emphasize that our observations do not exclude the possibility that MYPT1 plays a role in microtubule acetylation mediated regulation of myosin activity. Myosin activation is spatiotemporally modulated in directionally migrating cells. It is tempting to speculate that microtubule acetylation may control GEF-H1 and MYPT1 activity in distinct spatiotemporal manner to control myosin dynamics. Using optoTAT in these systems to test the impact of microtubule acetylation on myosin activation may help resolve these contradictory observations.

The critical role of GEF-H1 in optoTAT mediated myosin activation and rescue of chemotaxis defects in α-TAT1 KO MEFs by microtubule non-binding GEF-H1(C53R) (Fig. 6) suggest that GEF-H1 mediates crosstalk between actin and acetylated microtubules. Additionally, our data also suggest that GEF-H1 release is not mediated solely by microtubule stability, but through the recognition of the acetyl moiety, directly or indirectly (Fig. 5, Supplementary Fig. S4). How GEF-H1 can detect acetylated versus non-acetylated microtubules is an intriguing question. Since acetylation occurs in the microtubule lumen^6,9^, one possibility is that GEF-H1 enters the lumen to read the acetylation state of microtubules. Although GEF-H1 (∼100 kDa) is not a small molecule and its access to the narrow 15 nm diameter microtubule lumen appears difficult, larger molecules such as CSPP1 (∼138 kDa) have been reported to exist in the microtubule lumen^78^. The rapid release of GEF-H1 from microtubules, in such a case, would imply a somewhat permissive structure of a subset of microtubules, allowing for molecular exchange between the lumen and cytoplasm. Another possibility is that while GEF-H1 binds to the microtubule surface, it contains a domain which probes the lumen to detect the acetyl moiety, or the conformational changes in α-tubulin due to acetylation. It is also possible that GEF-H1 localization on microtubule is controlled by a third-party molecule that directly senses the acetylation state of microtubules, and relays that information to GEF-H1^70,79^.

How, or even if, microtubule acetylation mediates spatiotemporal control of GEF-H1 activity is an intriguing question. In order to achieve directional persistence in migrating cells, RhoA and myosin activation must be spatiotemporally regulated at sites of dynamic actin remodeling^80^, and so it may be reasoned that GEF-H1 activation is also spatiotemporally regulated. One possibility is that microtubule acetylation only releases GEF-H1 to increase the cytosolic pool, where GEF-H1 may be subcellularly activated through additional signaling pathways. Our data show comparable kinetics of optoTAT mediated microtubule acetylation, GEF-H1 release, myosin activation and adhesion maturation (Figs. 3, 4, 5), suggesting a direct and causal coupling of these events. On the other hand, α-TAT1 KO MEFs expressing GEF-H1(C53R) are capable of spatial regulation of myosin activation as well as directional migration (Fig. 6). This would suggest that beyond release from sequestration, microtubule acetylation may not spatially regulate GEF-H1 activation. Of course, we cannot rule out cellular adaptation in this instance. OptoTAT design does not allow subcellular activation of microtubule acetylation, thus limiting our capability of probing the effects of spatially restricted microtubule acetylation on cell behavior. α-TAT1 localizes to focal adhesions through Talin binding^22^, which may provide localized interaction with microtubules to facilitate spatially regulated GEF-H1 release and activation. Spatial distribution of GEF-H1 may be fine-tuned by combination of microtubule assembly-disassembly and acetylation state. Further examination of the spatial regulation of microtubule dynamics, microtubule acetylation and GEF-H1 activation may help us understand their interplay in migrating cells.

## Materials and Methods

### Cell culture and transfection

HeLa and HEK-293T cells were cultured in DMEM basal media and passaged every third day of culture. For optimal growth, the media were supplemented with 10% (v/v) fetal bovine serum, L-Glutamine, Penicillin/Streptomycin, Non-essential amino acids and 0.05 mM β-mercaptoethanol. WT and α-TAT1 KO MEFs were a generous gift from Dr. Maxence Nachury and were cultured in DMEM basal media supplemented with 10% (v/v) fetal bovine serum, L-Glutamine, Penicillin/Streptomycin, Non-essential amino acids and 0.05 mM β-mercaptoethanol. HeLa cells, WT MEFs and α-TAT1 KO MEFs stably transduced with mVenus-α-TAT1, mVenus-α-TAT1(D157N), mCherry-MRLC, mVenus-optoTATV2 or mCherry-GEF-H1(C53R) were sorted using the Sony SH800 cell sorter using manufacturer’s instructions to select cell populations with similar mVenus or mCherry fluorescence thresholds to ensure similar expression levels of the proteins of interest. The cells were maintained under standard cell culture conditions (37 °C and 5% CO_2_) and were checked for mycoplasma contamination prior to use in experiments. The stably transduced cells were cultured in medium containing 1 µg/ml of puromycin. Effective puromycin dosage was ascertained by testing on WT and α-TAT1 KO MEFs. FuGENE 6 reagent (Promega, Madison, WI) was used for transient transfection of HeLa cells according to the manufacturer’s instructions. For generation of lentiviral particles, HEK cells were transfected using polyethyleneimine (PEI). Electroporation with Lonza electroporator was performed for expression of VinTS in WT and α-TAT1 KO MEFs, according to the manufacturer’s instructions.

### DNA plasmids

α-TAT1 plasmid construct was a gift from Dr. Antonina Roll-Mecak. VinTS was a gift from Dr. Martin Schwartz (Addgene plasmid # 26019). mCherry-MRLC and mCherry-Paxillin were a gift from Dr. Yi I. Wu. NLS-mCherry-LEXY was a gift from Dr. Barbara Di Ventura & Dr. Roland Eils (Addgene plasmid # 72655). Z-lock αTAT was a gift from Dr. Klaus Hahn (Addgene plasmid # 175290). GFP-GEF-H1 was a gift from Dr. Hiroaki Miki. As indicated in the results and figure legends, tags of compatible fluorescent proteins including Cerulean, mVenus, mCherry and miRFP703 were appended to facilitate detection of the proteins of interest and the plasmids were subcloned into C1 vector (Clontech) or pTriEx4 vector (Novagen). Unless specified otherwise, the termini of tagging were positioned as in the orders they are written. Lentiviral plasmids were generated based on a modified Puro-Cre vector (Addgene plasmid # 17408, mCMV promoter and no Cre encoding region). Point mutations or truncations of indicated plasmid constructs were generated by PCR. The open reading frames of all DNA plasmids were verified by Sanger sequencing.

### Drug treatments

Pharmacological drugs were purchased as indicated: Y-27632 (LC Laboratories, catalog # Y-5301), Tubacin (Selleck Chemicals, catalog # S2239), Taxol or Paclitaxel (Cell Signaling Technology, catalog # 9807S). Y-27632 was applied at 10 μM final concentration 30 min before fixing cells or initiation of microscopy. Tubacin was applied at 2 μM final concentration for 4 hours before initiation of microscopy. Taxol was applied at 100 nM final concentration overnight before fixing cells.

### Immunofluorescence assays

For immunostaining of acetylated α-Tubulin, total α-Tubulin or GEF-H1, cells were fixed using ice-cold methanol for 10 minutes, washed thrice with cold PBS, blocked with 2% BSA in PBS for one hour and then incubated overnight at 4°C with antibodies against α-tubulin (rat, MilliporeSigma, MAB1864), acetylated α-Tubulin (mouse, MilliporeSigma, T7451) or GEF-H1 (rabbit, ThermoFisher, PA5-32213). Next day, the samples were washed thrice with cold PBS and incubated with secondary antibodies (Invitrogen) for one hour at room temperature, after which they were washed thrice with PBS and images were captured by microscopy. For immunostaining of Vinculin, phospho-MRLC, and Myosin IIa, cells were fixed using freshly prepared 4% paraformaldehyde at room temperature for 10 minutes, washed twice with PBS, blocked and permeabilized in 1% BSA in PBS with 0.1% TritonX-100 at room temperature for an hour and then incubated with antibody against Vinculin (mouse, Sigma Aldrich, MAB3574), phospho-MRLC (rabbit, Cell Signaling Technology, 3671T), Myosin IIa (rabbit, Cell Signaling Technology, 3403T) in the above blocking buffer at room temperature for 1 hour, washed three times in PBS and incubated with secondary antibody, Phalloidin (ThermoFisher A22286) and DAPI (Cell Signaling Technology, 4083S). After that, they were washed three times in PBS and the images were captured by microscopy. For HeLa cells transiently transfected with mCherry-α-Tubulin or mCherry-α-Tubulin(K40A), fixing and immunostaining were performed 24 hours post-transfection. For Y-27632 treatment, HeLa cells were treated with 10 μM Y-27632 or equal volume of vehicle (water), incubated for 4 hours, followed by PFA fixation and immunostaining. For Taxol treatment, WT or α-TAT1 KO MEFs were treated with Taxol (100 nM) or vehicle (DMSO) overnight followed by methanol fixation and immunostaining.

### Western blot assays

Cell lysates were prepared by scraping cells using lysis buffer (RIPA buffer, Cell Signalling # 9806S), mixed with protease/phosphatase inhibitor cocktail (Cell Signaling # 5872S). Cell lysates were rotated on a wheel at 4°C for 15 min and centrifuged for 10 min at 15,000 g 4°C to pellet the cell debris, mixed with NuPAGE™ LDS Sample Buffer (Thermo Fisher # NP0007) with protease and phosphatase inhibitors and boiled 5 min at 95°C before loading in polyacrylamide gels. Gels were transferred and membranes were blocked with TBST (0.1% Tween) and 5% BSA and incubated overnight with the primary antibody, and 1 h with LiCOR IR-dye conjugated secondary antibodies after which bands were revealed using Odyssey imaging system. Antibodies used: anti-α-Tubulin (rat, MilliporeSigma, MAB1864), anti-Vinculin (rabbit, Cell Signaling Technology, 13901S), anti-GEF-H1 (rabbit, ThermoFisher, PA5-32213). Secondary IR-dye conjugated antibodies were purchased from LiCOR.

### Microscopy and image analyses

All epifluorescence imaging was performed with an Eclipse Ti microscope (Nikon) with PCO.Edge sCMOS camera (Excelitas), driven by NIS Elements software (Nikon). All TIRF imaging was performed with an Eclipse Ti microscope (Nikon) with ORCA-FusionBT sCMOS camera (Hamamatsu), driven by NIS Elements software (Nikon). All confocal images were captured using a laser scanning confocal Eclipse Ti2 microscope (Nikon) equipped with a tunable GaAsp detector and 2k resonant scanner (AXR, Nikon) with camera, driven by NIS software (Nikon). All live cell imaging was conducted at 37°C, 5% CO_2_ and 90% humidity with a stage top incubation system (Tokai Hit). Vitamin and phenol red-free media (US Biological) supplemented with 2% fetal bovine serum were used in imaging to reduce background and photobleaching. Inhibitors and vehicles were present in the imaging media during imaging. All image processing and analyses were performed using Metamorph (Molecular Devices, Sunnyvale, CA, USA) and FIJI software (NIH, Bethesda, MD, USA). OptoTAT stimulation was provided by epifluorescent 440 nm excitation 20 s apart, or continual exposure to blue LED light (Amazon, B08FQSFFJ60). Cells that were rounded up or showed a high degree of blebbing were excluded from analysis to minimize artifacts from mitotic, apoptotic or dead cells. For all ratiometric or intensity analyses, background subtraction based on a cell free area on each image was performed prior to calculation of the ratio. For colocalization analysis, Coloc2 function in FIJI was used to calculate the Pearson’s correlation coefficient (also called Pearson’s R) value. Images containing any saturated pixels in any channel (65535 value) within the cell area were excluded. The ratio of acetylated α-Tubulin over α-Tubulin (Ac. α-Tub/α-Tub) for transiently transfected cells was normalized against that for non-transfected cells averaging over >20 non-transfected cells from the same dish. For mVenus-optoTAT mediated kinetics of microtubule acetylation, only ratio of acetylated α-Tubulin over α-Tubulin (Ac. α-Tub/α-Tub) for individual cells are shown. Changes in mCherry-MRLC distribution isotropy was analyzed by OrientationJ plugin in FIJI^81^.

### Cell migration assays

Random migration assays were performed in 24-well plates. 1.5 x 10^4^ WT MEFs or 1.2 x 10^4^ α-TAT1 KO MEFs were seeded and incubated for 4-5 hours. After attachment, cells were imaged using phase contrast microscopy with 10X objective every 10 minutes for 15 hours. The wound healing assay was performed using Ibidi Culture-Insert 3 Well in 24-well plates (Ibidi, catalog # 80369). 3.5 x 10^4^ WT and α-TAT1 KO MEFs were seeded in each insert well and incubated for 4-5 hours or until a full monolayer was created. After cells were settled, the 3-well insert was removed, creating a 500 μm cell-free area between cell monolayers. Cells were imaged using phase contrast microscopy with 10X objective every 5 minutes for 15 hours. The wound closure rate was calculated by determining the area of the cell-free area over time. Chemotaxis assay was performed using μ-slide chemotaxis chambers coated with collagen IV (Ibidi, catalog # 80322) following the manufacturer’s protocol. In short, 2.5 x 10^6^ cells/mL αTAT1 KO MEFs, 3 x 10^6^ cells/mL WT MEFs or 3 x 10^6^ cells/mL KO MEFs rescued with mVenus-α-TAT1 or mCherry-GEF-H1 (C53R), were seeded with serum-free DMEM and incubated at 37°C, 95% humidity, and 5% CO2 for 4-5 hours. After cells were settled, serum-free DMEM was added to the right and left reservoirs of the chamber. Half of the volume of the left reservoir was replaced with DMEM supplemented with 20% FBS to generate the chemoattractant gradient. Cells were imaged using phase contrast with 10X objective every 10 minutes for 15 hours at 37°C and 5% CO2. Tracking and analyses were performed using ImageJ plug-ins, MTrackJ, and Chemotaxis_tool, respectively.

### Statistical analyses and reproducibility

Microsoft Excel (Microsoft, Redmond, WA, USA) and R (R Foundation for Statistical Computing, Vienna, Austria) were used for statistical analyses. The exact number of samples for each data set is specified in the respective figure legends. For live cell experiments, data were pooled from at least three independent experiments performed on different days. For immunocytochemistry data, compared data were collected from experiments performed in parallel with cells plated on the same 8 well chambers (Cellvis, C8-1.5H-N) on the same day with the same reagents and imaging performed under the same conditions. Individual cells were identified based on Phalloidin or α-Tubulin staining. Sample sizes were chosen based on the commonly used range in the field without performing any statistical power analysis and assumed to follow normal distribution. *P*-values were obtained from two-tailed Student’s *t*-test assuming equal variance or paired *t-*test where applicable.

## Acknowledgements

We thank Dr. Allen Kim for discussion that led to initiation of this project. We thank Dr. Maxence V. Nachury for WT and α-TAT1 KO MEF cells. We thank Dr. Sandrine Etienne-Manneville, Dr. Shailaja Seetharaman and Dr. Anna Akhmanova for insightful comments on the project. We thank Robert DeRose for manuscript proofreading and experimental support.

## Author contributions

ADR initiated the project. ADR and TI designed the experiments. ADR, CSG, FS, EY performed the experiments and data analyses under guidance from TI. ADR wrote the manuscript in consultation with CSG, FS and EY. TI edited the manuscript. All authors contributed to the final version of the manuscript.

## Data availability

All data and plasmid constructs will be made available on reasonable requests.

## Supplementary information

**Supplementary Figure S1.**
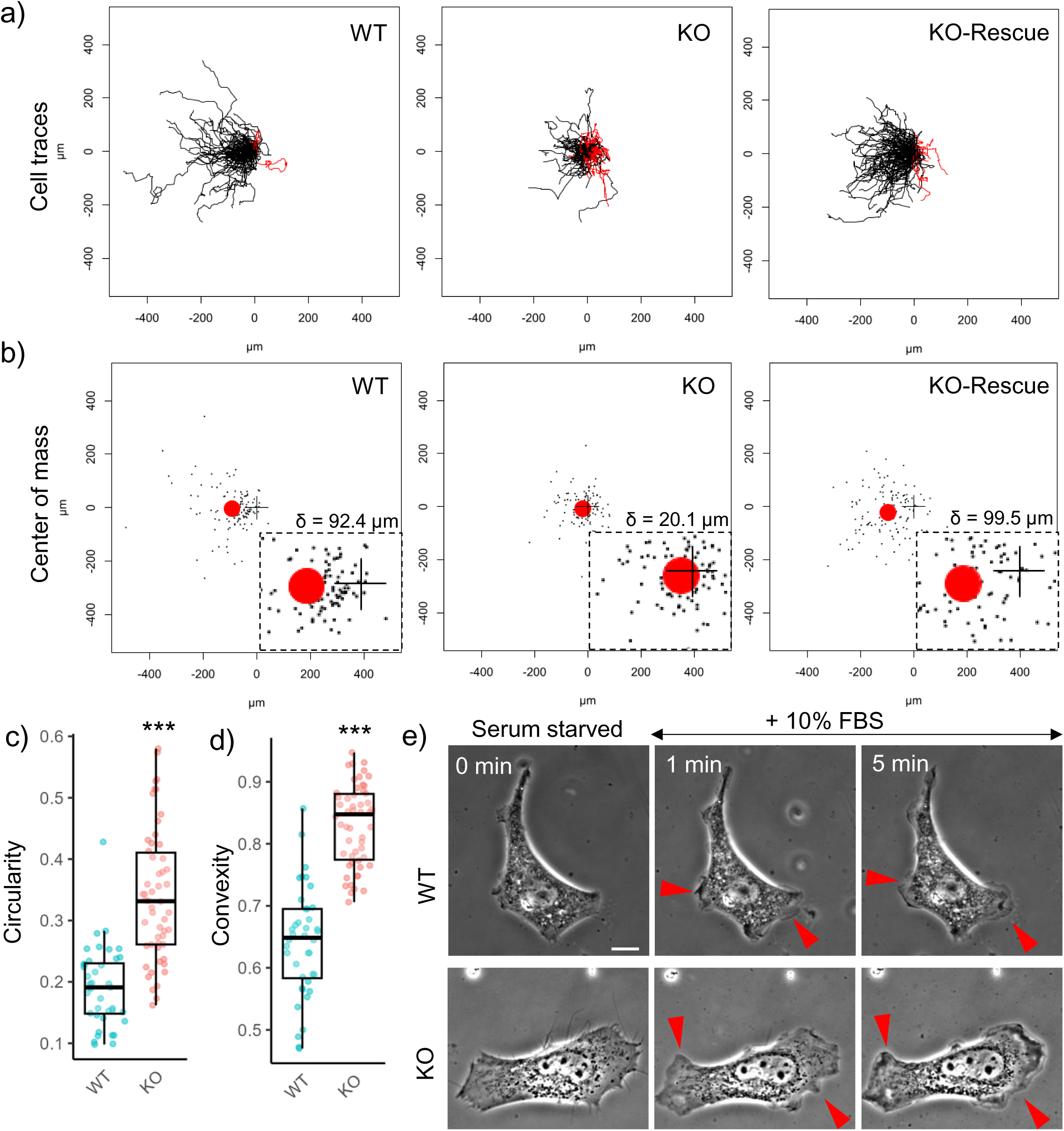
a) Tracks of WT, α-TAT1 KO and KO-rescue with α-TAT1 MEFs in chemotaxis assay, n = 120 cells (40 each from three independent experiments); b) Final location of individual cells (black dots) and the center of mass of all the cells (red circle) in chemotaxis assay, origin is indicated by “+”, distance between origin and center of mass (δ) is shown above the inset; c) Circularity and d) Convexity of WT α-TAT1 KO MEFs (WT: 40 and KO: 54 cells); e) Morphological changes in serum-starved WT and α-TAT1 KO MEFs on addition of 10% FBS, induced protrusions are indicated with red arrowheads, scale bar: 10 μm. ***: p<0.001

**Supplementary Figure S2.**
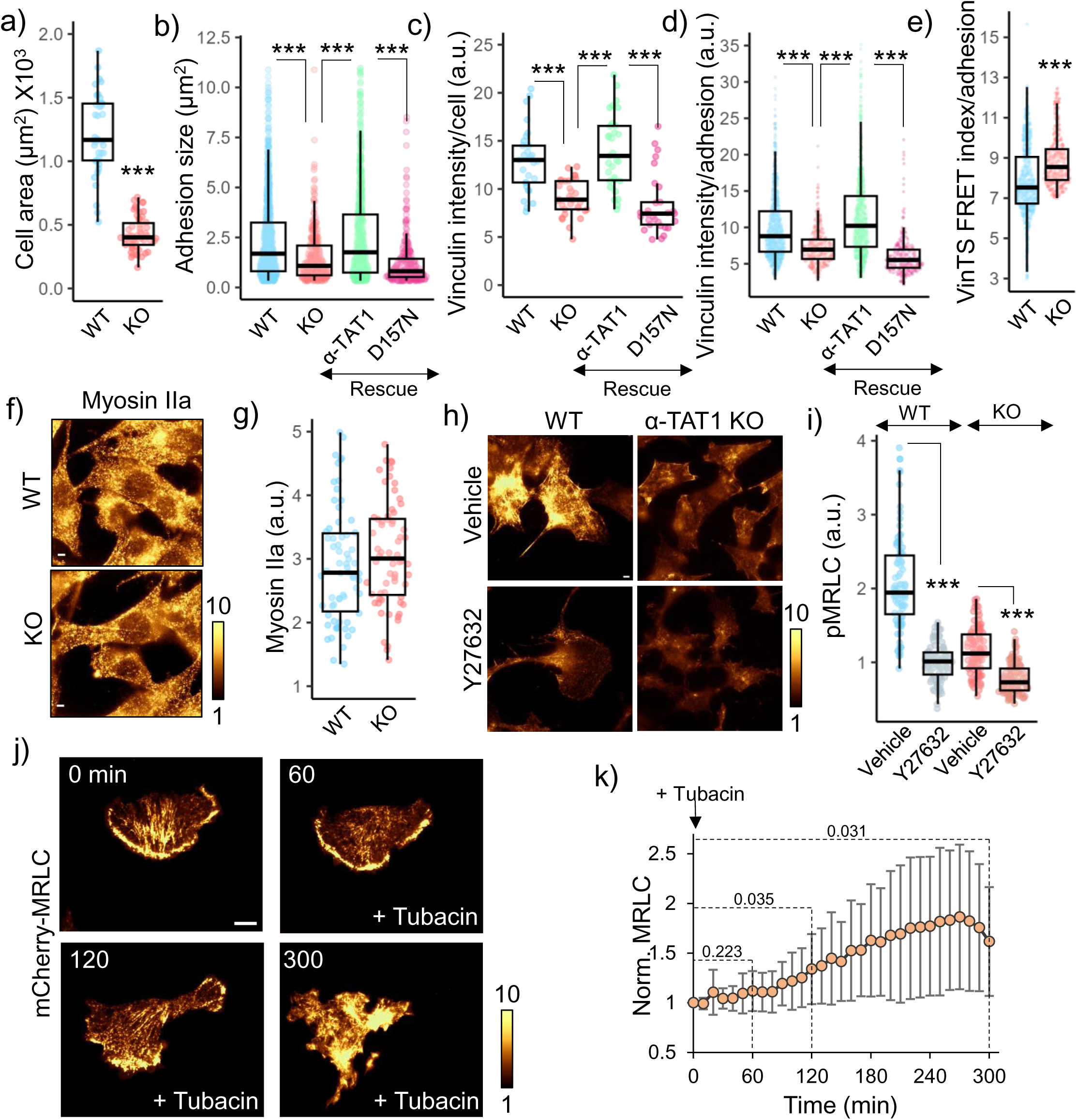
a) Cell areas of WT and α-TAT1 KO MEFs (WT: 40, KO: 54 cells); b) adhesion sizes, c) average vinculin intensity per cell, d) average vinculin intensity per adhesion in WT, α-TAT1 KO, rescue-WT and rescue-D157N MEFs (WT: 20, KO: 17, rescue-WT: 16, rescue-D157N: 22 cells); e) VinTS FRET index per adhesion in WT and α-TAT1 KO MEFs (WT:: 18, KO: 16 cells); f), g) Myosin IIa levels in WT and α-TAT1 KO MEFs (WT: 69, KO: 65 cells); h), i) Phospho-MRLC levels in WT and α-TAT1 KO MEFs treated with vehicle or 10 µM Y-27632 (WT-vehicle: 88, WT-Y27632: 91, KO-vehicle: 89, KO-Y27632: 98 cells); j) TIRF images of and k) changes in fluorescence intensity of mCherry-MRLC in WT MEFs on tubacin treatment, 12 cells, mean ± 95% C.I.; scale bar: 10 μm. ***: p<0.001

**Supplementary Figure S3.**
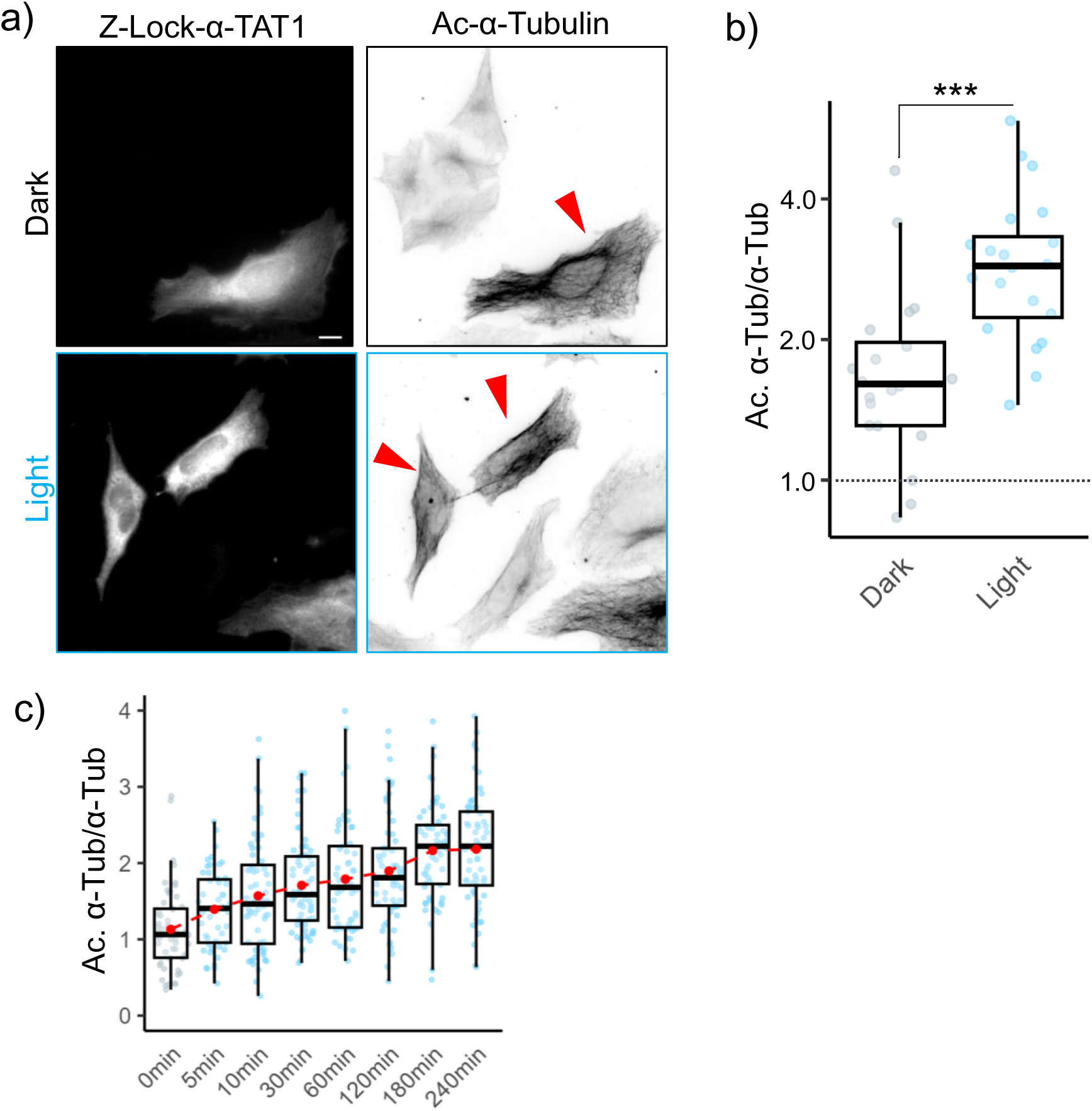
a) Microtubule acetylation levels in HeLa cells exogenously expressing mCherry-Z-Lock-α-TAT1, kept in dark or exposed to blue light for 2 hours, red arrowheads indicate transfected cells; scale bar: 10 μm; b) Microtubule acetylation levels in Hela cells expressing mCherry-Z-Lock-α-TAT1 in dark or after blue light exposure, normalized against acetylation levels in non-transfected cells; c) Temporal changes in acetylated microtubules (normalized against total α-Tubulin) on blue light stimulation of HeLa cells stably expressing mVenus-optoTAT V2 (0 min: 54, 5 min: 50, 10 min: 61, 30 min: 66, 60 min: 61, 120 min: 62, 180 min: 60 and 240 min: 61 cells), red dots indicate the mean values; note: time scale is not linear.

**Supplementary Figure S4.**
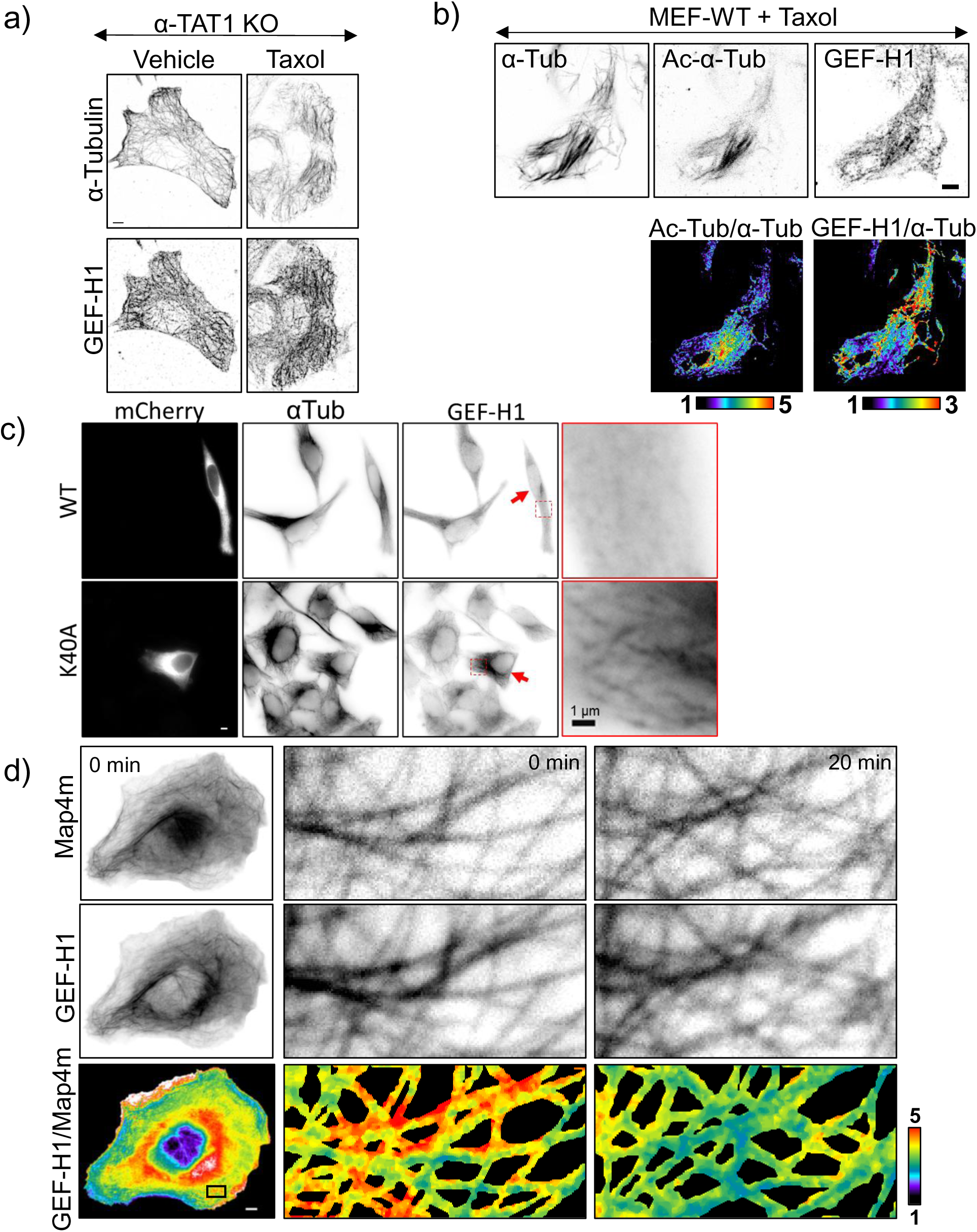
a) Immunostaining against α-Tubulin and GEF-H1 α-TAT1 KO MEFs treated with vehicle (DMSO) or 100 nM Taxol overnight; b) Immunostaining against α-Tubulin, acetylated α-Tubulin and GEF-H1 in WT MEFs treated with 100 nM Taxol overnight; c) Immunostaining against α-Tubulin and GEF-H1 in HeLa cells expressing mCherry-α-Tubulin or mCherry-α-Tubulin(K40A) (lower panels), transfected cells are indicated with red arrowheads, insets are magnified on the right panel; d) Changes in mCherry-GEF-H1/mVenus-MAP4m signal in HeLa cells expressing miRFP703-optoTAT on blue light stimulation, inset is magnified in the right panels; Scale bar: 10 μm or as indicated.

**Supplementary Figure S5.**
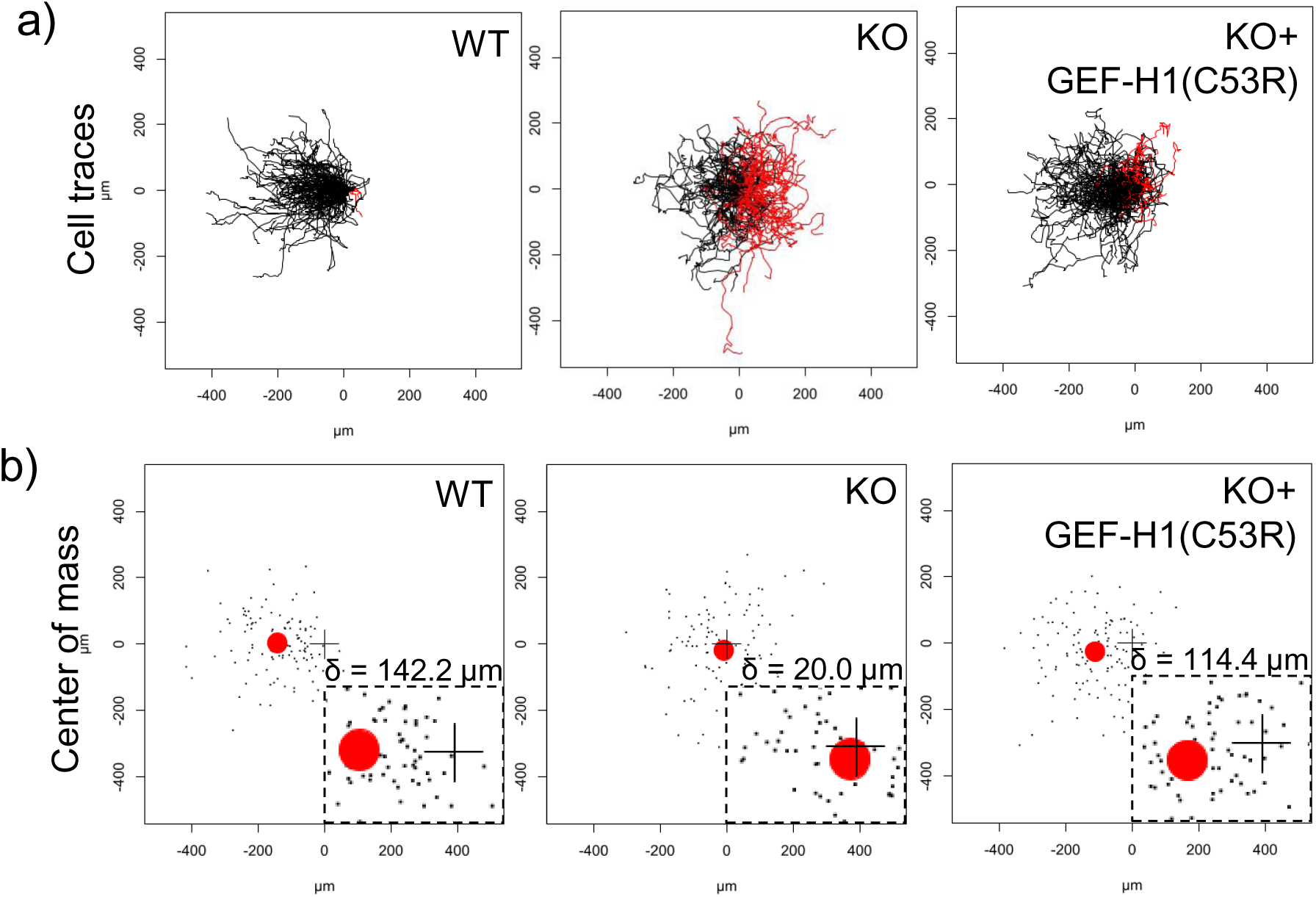
a) Tracks of WT, α-TAT1 KO and KO-rescue with mCherry-GEF-H1(C53R) MEFs in chemotaxis assay, n = 120 cells (40 each from three independent experiments); b) Final location of individual cells (black dots) and the center of mass of all the cells (red circle) in chemotaxis assay, origin is indicated by “+”, distance between origin and center of mass (δ) is shown above the inset.

